# Accurate and General DNA Representations Emerge from Genome Foundation Models at Scale

**DOI:** 10.1101/2024.12.01.625444

**Authors:** Caleb N. Ellington, Ning Sun, Nicholas Ho, Tianhua Tao, Sazan Mahbub, Dian Li, Yonghao Zhuang, Hongyi Wang, Le Song, Eric P. Xing

## Abstract

Language models applied to protein sequences have become a panacea, enabling therapeutics development, materials engineering, and core biology research. Despite the successes of protein language models, genome language models remain nascent. Recent studies suggest the bottleneck is data volume or modeling context size, since long-range interactions are widely acknowledged but sparsely annotated. However, it may be the case that even short DNA sequences are modeled poorly by existing approaches, and current models are unable to represent the wide array of functions encoded by DNA. To study this, we develop AIDO.DNA, a pretrained module for DNA representation in an AI-driven Digital Organism [1]. AIDO.DNA is a seven billion parameter encoder-only transformer trained on 10.6 billion nucleotides from a dataset of 796 species. By scaling model size while maintaining a short context length of 4k nucleotides, AIDO.DNA shows substantial improvements across a breadth of supervised, generative, and zero-shot tasks relevant to functional genomics, synthetic biology, and drug development. Notably, AIDO.DNA outperforms prior encoder-only architectures *without* new data, suggesting that new scaling laws are needed to achieve compute-optimal DNA language models. Models and code are available through Model-Generator in https://github.com/genbio-ai/AIDO and on Hugging Face at https://huggingface.co/genbio-ai.

## 1 Introduction

Genomes are the product of billions of years of evolution and selection under the forces of fitness, competition, niches, and stochasticity. They are a nearly universal requirement for life, encoding all cellular systems and their components – RNA, proteins, their assembly, and their regulation – through a simple 4-chemical vocabulary allowing self-replication, recombination, and inheritance. The conservation of genomic elements across evolutionary timelines is highly correlated to their functional roles and interactions within the living systems they encode. Aligning sequences to determine their elementwise conservation has helped to discover protein structures, binding sites, regulatory elements, 3D genome structure, pathologies, and genealogies. However, function is difficult to determine from conservation alone. Instead, conservation analysis often plays the role of data preprocessing for feature selection, which has limited downstream analyses to use features which are well-sampled, well-characterized, and conserved across species. New computational and statistical tools are required to go beyond this regime.

Recently, large unsupervised pretrained foundational models for predicting protein sequence conservation have achieved widespread success on difficult and diverse tasks that decades of prior work has sought to address, including protein folding and *de novo* design [19, 20], exonic variant pathogenicity prediction [21, 22], and representation of evolutionary dynamics beyond multiple sequence alignment [23]. These conservation models accurately predict protein structure, catalytic activity, and a protein’s role and fitness within a broader biological system, the same way a traditional multiple-sequence alignment would be used, while also generalizing to human-only and novel sequences [21], and generating new sequences for desired functions [24]. Despite these successes, the architectures for protein foundation models have not shown the same success when applied more broadly to genomes [25, 26, 27].

Recent works suggest the gap in performance may be related to the length of the total genomic context accommodated by the model, and thus propose new architectures to learn long-range interactions and their role in genetic regulation [8, 13, 10]. However, increasing context length to millions of nucleotides has shown diminishing returns [9]. While the field appears at an impasse, reviewing the growth of deep learning models in genomics reveals successful variants and their proliferation in research (Table 1). While no consensus has been reached on the optimal context size for representation learning, parameter sizes continue to grow and vanilla BERT-style encoder-only transformer architectures have been widely successful.

**Table 1:**
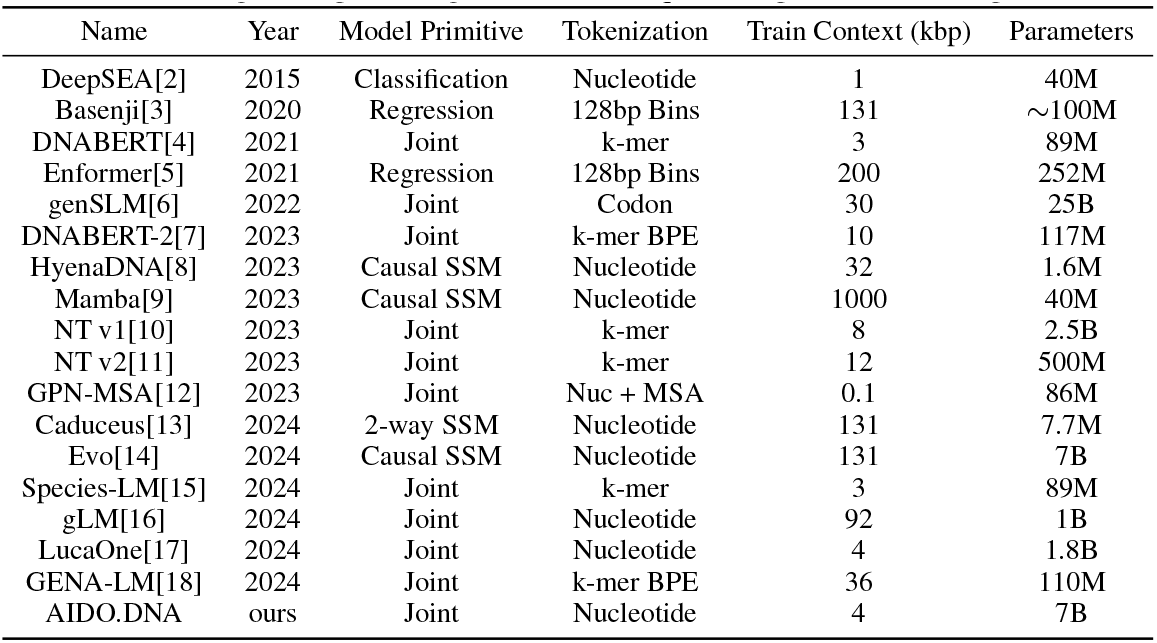
Catalog of deep learning methods for sequence representation and prediction.

However, DNA encoders are still dwarfed by their protein cousins [24]. This is alarming, since a general genome language model should implicitly contain a protein language model while also representing all other molecular functions encoded by DNA. Given this broad modeling scope, we believe that these models require vastly more representation capacity than has previously been attempted. To learn accurate and general representations of genomic functions, we develop AIDO.DNA, one of the largest unsupervised pretrained DNA encoders to date at seven billion parameter scale. We train this model at single nucleotide resolution on 796 species’ genomes with 10.6 billion nucleotides. To address concerns about context length, we explicitly set the context to a relatively short 4,000 nucleotides. By achieving a new state-of-the-art (SOTA) on canonical tasks in transfer learning, zero-shot prediction, and sequence generation, AIDO.DNA represents an important step toward the development of an AI-driven Digital Organism [1], and directly enables new approaches to core biology research, genomics, synthetic biology, metabolic engineering, and therapeutics design.

## 2 Methods

We develop AIDO.DNA at 300M and 7B parameter scales, using large-scale distributed pretraining [28] and parameter efficient transfer learning frameworks [29] to rapidly scale pretraining and finetuning. Our model is based on the bidirectional transformer encoder (BERT) architecture [30] with single-nucleotide tokenization, and is optimized using a masked language modeling (MLM) training objective.

### 2.1 Pretraining

#### Data

To test whether representation capacity has limited the development of DNA language models in previous studies, we utilize the data set and splits from the Nucleotide Transformer [10]. Starting from a total of 812 genomes with 712 for training, 50 for validation, and 50 for testing, we removed 17 entries which had been deleted from NCBI since the original dataset’s publication on Hugging Face. One of these was the important model organism *Rattus norvegicus*, which we replaced with the current reference genome. This resulted in 696 genomes for training, 50 for validation, and 50 for testing. With a total of 10.6 billion training tokens, we pretrained AIDO.DNA at 300M and 7B parameter scales.

#### Tokenization

To appropriately adapt langauge models to biology, it is important to be clear that DNA is not a sentence. DNA is an aperiodic crystal composed of four chemical building blocks – the nucleotides adenine (A), thymine (T), cytosine (C), and guanine (G) – which are paired and polymerized on a phosphate backbone. DNA has no periods, no sentences, no paragraphs, and no global grammar. With its very small vocabulary, DNA encodes an enormous diversity of biomolecules and functions. Compared with natural language, it has a very low per-character information density on average, and grammars are highly local, relativistic, and context-specific. Prior works which compress nucleotides into “words” [10, 7] make it difficult to learn relative and local grammars (e.g. codons) and dilute the representation of high impact context-specific effects such as single nucleotide pathogenic variants [31].

To minimize bias and learn high-resolution single-nucleotide dependencies, we opted to align closely with the real data and use character-level tokenization with a 5-letter vocabulary: A, T, C, G, N, where N is commonly used in gene sequencing to denote uncertain elements. Sequences were also prefixed with a [CLS] token and suffixed with a [EOS] token as hooks for downstream tasks. We chose a context length of 4,000 nucleotides as the longest context which would fit within our largest 7B model size during pretraining, and chunked our dataset of 796 genomes into non-overlapping segments.

#### Architecture

To learn semantically meaningful representations, we employed an BERT-style encoder-only dense transformer architecture [30]. We make minor updates to this architecture to align with current best practices, including using SwiGLU [32] and LayerNorms [33]. Additionally, we use Rotary Positional Embeddings (RoPE) [34], given that DNA syntax does not function based on absolute nucleotide positions but nucleotides interact in highly local and context-specific ways. Our 300M model used 24 transformer blocks with embedding size 1,024, where each block contains 16 self-attention heads and a feed forward size 2,688. Our 7B model used 32 transformer blocks with embedding size 4,352, where each block contains 32 self-attention heads and feed forward size of 11,584 (Figure 1).

**Figure 1.**
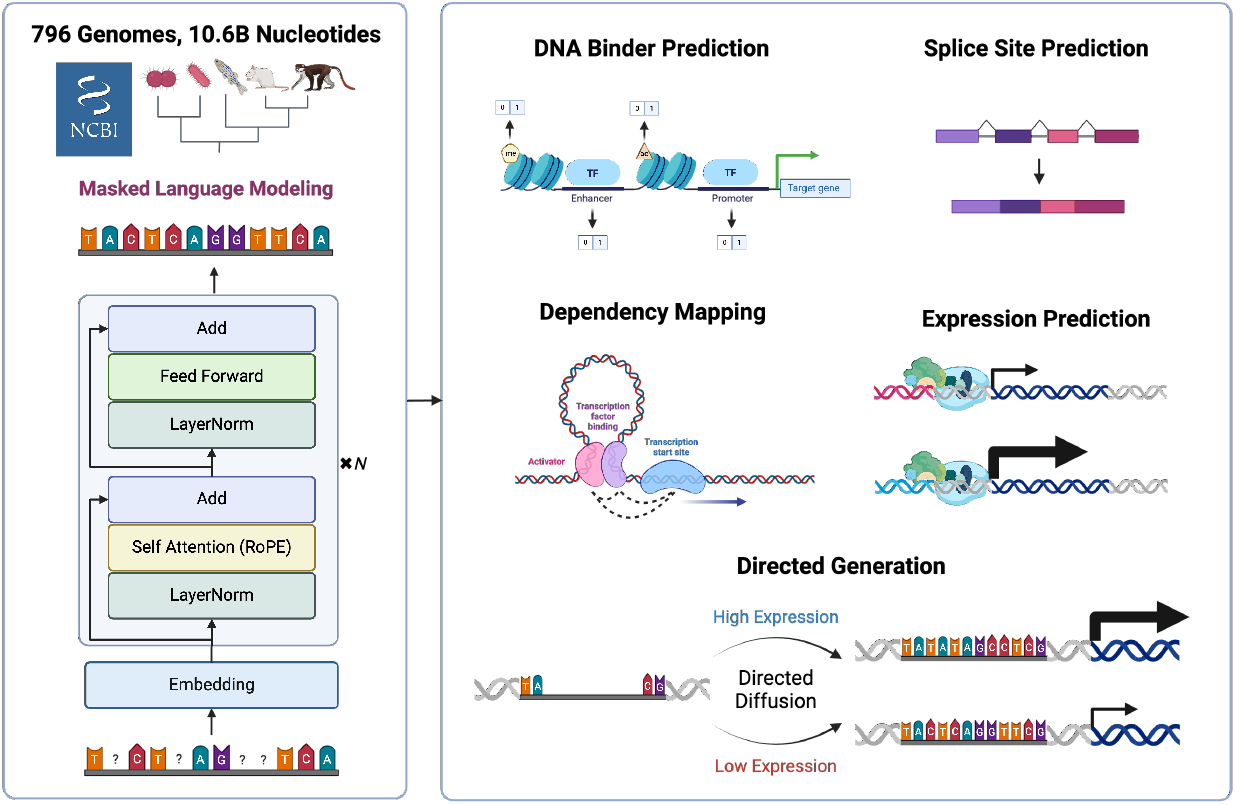
AIDO.DNA pretraining (left) and evaluation (right).

#### Training

The weights of our seven billion parameter model occupy over 200GB of memory in 32 bit precision. To train a model of this size, we use model parallelism to split training across 256 H100 GPUs using the Megatron LM framework [28]. We also employed bfloat16 mixed precision training and FlashAttention-2 [35] to allow for training with large context length at scale. With these optimizations, our maximum possible global batch size was 1024 with a per-device micro batch size of 2. With this configuration, the model took 8 days to train to 100,000 iters. The 300 million parameter model trained for 4 days on 32 A100 GPUs. Full training configurations and practical considerations for both models are available in the Appendix D.1.

#### Optimization

We trained AIDO.DNA using a classic masked language modeling (MLM) objective, choosing 15% of tokens in each sequence at random to alter. Of the chosen tokens, 80% are masked, 10% are corrupted (uniformly re-sampled), and 10% are untouched. We use a cross-entropy loss on these tokens, comparing with the original unaltered sequence. We used the Adam optimizer with *β*_1_ = 0.9 and *β*_2_ = 0.95. We also used a cosine learning rate scheduler with a 2% linear warmup and a minimum learning rate of 10^*−*5^.

### 2.2 Evaluation

We evaluate the benefits of pretraining our 300M and 7B models by conducting a comprehensive series of experiments related to functional genomics, genome mining, metabolic engineering, synthetic biology, and therapeutics design, covering supervised, unsupervised, and generative objectives. Unless otherwise stated, hyperparameters were determined by optimizing model performance on a 10% validation split of the training data, and models were tested using the checkpoint with the lowest validation loss.

#### Sequence Property Classification

We apply the pretrained AIDO.DNA models to sequence classification and property prediction tasks, using standard classification benchmarks from prominent works on DNA encoders covering a breadth of genomic functions related to transcriptional regulation and transcript processing [10, 7]. We use a binary cross-entropy objective, full parameter finetuning with [CLS] pooling and a linear prediction head. For transcription factor classification tasks, we continued our unsupervised masked language modeling pretraining objective for several epochs on test set input sequences before applying the supervised finetuning objective, given recent work on the importance of well-aligned inductive biases in dense transformers [36].

#### Zero-shot Variant Effect Prediction

To evaluate the utility of AIDO.DNA’s pretrained embeddings in a zero-shot setting, we use a curated set of ClinVar pathogenic and GnomAD benign variant data to predict variant effects [10]. We inferred embeddings for each sequence variant by performing inference on a 1kbp window centered on the variant, then averaging embeddings across the sequence length. Assuming benign variants will produce similar embeddings to the reference while pathogenic variants will produce different embeddings, we quantify the AUROC of our model by taking the max-normalized L2 distances between reference and variant sequence embeddings as the probability of variant pathogenicity.

#### Dependency Mapping for Genome Mining

We apply AIDO.DNA to discover unnannotated but functionally important regions of the human genome by implementing an *in silico* mutagenesis-based nucleotide dependency mapping strategy [31]. This method produces a 2D grid of dependency values between each nucleotide, indicating their co-conservation and likely functional dependency. The grid of predicted dependencies is composed of dependency pixels *e*_*i,j*_ for each nucleotide pair *i* and *j*, where

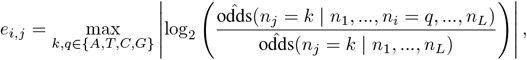

such that *k* and *q* are the key and query nucleotide types, *n* is a length *L* DNA sequence, and 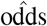 are the odds inferred using the pretrained AIDO.DNA. Crucially, this method leverages the pretrained model without finetuning, allowing us to evaluate the richness of representations inferred from unsupervised learning alone.

#### Predicting Gene Expression from Genomic Contexts

We propose a new variant of the canonical task of predicting gene expression from genomic contexts based on a dataset of 10 million synthetic promoter sequences with paired expression measurements [37]. This dataset enjoys desirable properties for downstream analysis and evaluation of models, well-characterized transcription factor-promoter interactions and random sequence construction, which affords useful analytical properties. We finetune AIDO.DNA using the same setup as classification with a mean squared-error objective to predict gene expression from sequence.

#### Directed Promoter Generation

Applying AIDO.DNA to generate functional genetic code, we invert our gene expression prediction task to generate promoter sequences toward a desired expression level. MLM-trained models are not suitable for complex sequence generation, so we finetune AIDO.DNA with a simple masked diffusion language modeling objective [38]. To add the expression level condition, we append our pretrained model with 2 transformer blocks, whose input is the pretrained embeddings added to a linear expression level encoding. To allow our model to ignore conditioning information if it is uninformative of the data distribution and leverage the pretrained decoder head, we also add a skip connection from the pretrained embeddings to the output of the conditioned transformer blocks.

## 3 Results

We developed AIDO.DNA, the largest and most performant encoder-only foundation model to date for representation, transfer learning, and generation of DNA sequences. While long context models have dominated recent literature, AIDO.DNA shows that substantial gains can be made on most tasks by scaling model depth on a short context length of 4,000 tokens at single-nucleotide resolution. On suites of transfer learning benchmarks from recent works, AIDO.DNA achieves a new state-of-the-art on sequence property prediction and zero-shot variant effect prediction (Fig. 2). Furthermore, pretraining at scale reveals functional dependencies without the need for curated finetuning data, helping to define and annotate new regulatory elements (Fig. 3). Finally, we show that AIDO.DNA enables directed design of promoter sequences toward a desired expression level (Table 6).

**Figure 2.**
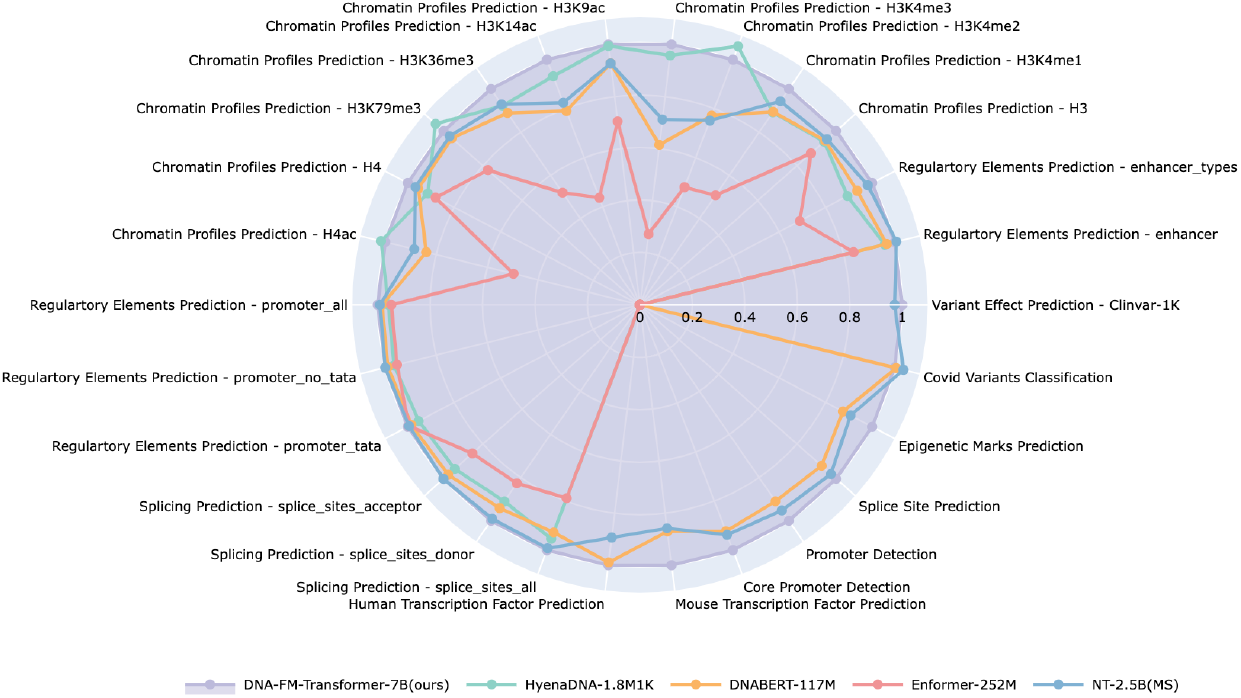
Benchmarking summary for AIDO.DNA, normalized to AIDO.DNA performance.

**Figure 3.**
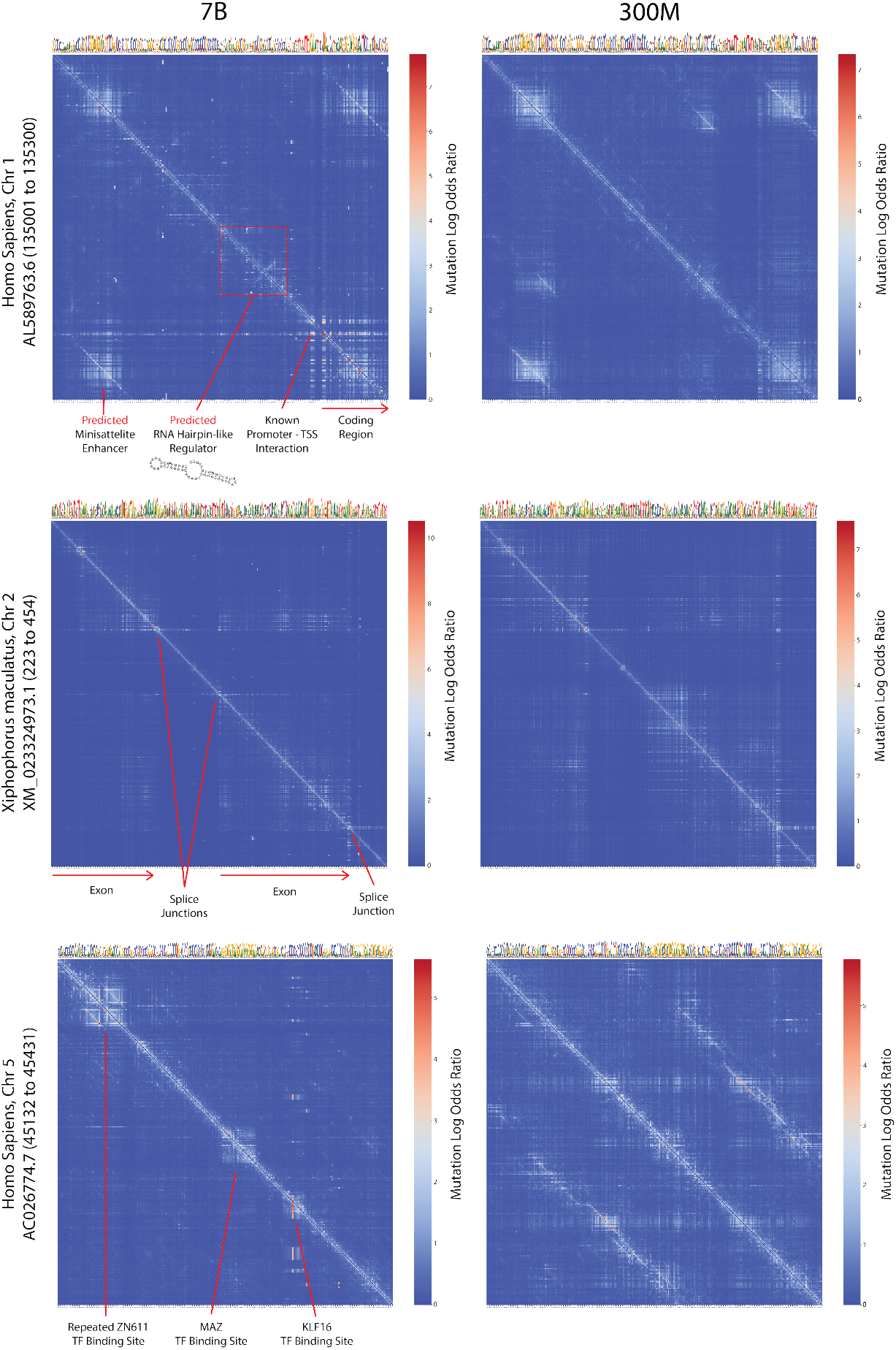
Unsupervised discovery of functional genomic elements using *in silico* mutagenesis. From pretraining solely on DNA, AIDO.DNA learns to identify RNA, protein, and regulatory elements encoded in DNA, as well as their interactions. Scaling to 7B parameters reveals highly specific and fine-grained interactions ignored at smaller scales.

### 3.1 Scaling DNA Encoders to Seven Billion Parameters

AIDO.DNA is the largest encoder-only DNA foundation model to date, with 7 billion parameters trained on 10.6 billion nucleotides from a multi-species dataset of 796 species. This multispecies dataset is nearly identical to the one used by Nucleotide Transformer [10], after removing genomes which have since been deleted from NCBI. In comparison with other encoder-only models Nucleotide Transformer [10] and DNABERT [4] models, AIDO.DNA also makes only modest architecture changes. However, we find that the main limitation of such models is their size. By scaling DNA encoders to seven billion parameters, we see substantial gains across standard benchmarks, and notice a variety of improvements to qualitative tasks relevant to genome mining, metabolic engineering, and synthetic biology (Tables 2, 3, 4, 5, 6).

**Table 2:** Genome Understanding Evaluation Benchmarks [7]. Metrics are Matthew’s Correlation Coefficient (MCC) unless otherwise stated. DB is DNABERT. NT is Nucleotide Transformer. ^*†*^Uses self pre-training before supervised finetuning [36]

| Task | DB<br>89M | DB-2 old<br>117M | DB-2<br>117M | NT<br>2.5B | AIDO.DNA<br>300M | AIDO.DNA<br>7B |
| --- | --- | --- | --- | --- | --- | --- |
| human-TF-0 | 66.84 | <u>71.99</u> | 69.12 | 66.64 | 68.192 | **72.216**<sup>†</sup> |
| human-TF-1 | 70.14 | <u>76.06</u> | 71.87 | 70.28 | 70.656 | **77.929**<sup>†</sup> |
| human-TF-2 | 61.03 | 66.52 | 62.96 | 58.72 | <u>70.987</u> | **72.273**<sup>†</sup> |
| human-TF-3 | 51.89 | **58.54** | <u>55.35</u> | 51.65 | 47.214 | 55.129<sup>†</sup> |
| human-TF-4 | 70.97 | **77.43** | 74.94 | 69.43 | 75.003 | <u>76.879</u><sup>†</sup> |
| mouse-TF-0 | 44.42 | 56.76 | <u>64.23</u> | 63.31 | 56.71 | **67.292**<sup>†</sup> |
| mouse-TF-1 | 78.94 | 84.77 | <u>86.28</u> | 83.76 | 82.447 | **86.295**<sup>†</sup> |
| mouse-TF-2 | 71.44 | 79.32 | 81.28 | 71.52 | <u>84.729</u> | **91.110**<sup>†</sup> |
| mouse-TF-3 | 44.89 | 66.47 | 73.49 | 69.44 | <u>77.246</u> | **86.769**<sup>†</sup> |
| mouse-TF-4 | 42.48 | <u>52.66</u> | 50.8 | 47.07 | 47.848 | **55.375**<sup>†</sup> |
| core_promoter_all | 68.9 | 69.37 | 67.5 | 70.33 | **72.977** | 71.80 |
| core_promoter_no_tata | 70.47 | 68.04 | 69.53 | 71.58 | **72.054** | <u>71.623</u> |
| core_promoter_tata | 76.06 | 74.17 | 76.18 | 72.97 | <u>82.221</u> | **85.045** |
| Promoter all | 90.48 | 86.77 | 88.31 | 91.01 | <u>93.229</u> | **93.532** |
| promoter no TATA | 93.05 | <u>94.27</u> | 94.34 | 94 | 94.165 | **94.955** |
| Promoter TATA | 61.56 | 71.59 | 68.79 | 79.43 | <u>86.057</u> | **89.104** |
| Splice Reconstruction | 84.07 | 84.99 | 85.93 | 89.35 | <u>90.727</u> | **91.16** |
| H3 | 73.1 | 78.27 | <u>80.17</u> | 78.77 | / | **82.485** |
| H3K14ac | 40.06 | 52.57 | <u>57.42</u> | 56.2 | / | **64.927** |
| H3K36me3 | 47.25 | 56.88 | 61.9 | <u>61.99</u> | / | **67.441** |
| H3K4me1 | 41.44 | 50.52 | 53 | <u>55.3</u> | / | **57.826** |
| H3K4me2 | 32.27 | 31.13 | <u>39.89</u> | 36.49 | / | **47.031** |
| H3K4me3 | 27.81 | 36.27 | <u>41.2</u> | 40.34 | / | **44.788** |
| H3K79me3 | 61.17 | <u>67.39</u> | 65.46 | 64.7 | / | **68.278** |
| H3K9ac | 51.22 | 55.63 | <u>57.07</u> | 56.01 | / | **64.17** |
| H4 | 79.26 | 80.71 | <u>81.86</u> | 81.67 | / | **81.947** |
| H4ac | 37.43 | <u>50.43</u> | 50.35 | 49.13 | / | **61.2** |
| COVID (F1) | 55.5 | <u>71.02</u> | 68.49 | **73.04** | 69.714 | 70.575 |
| Average TF | 64.174 | <u>70.108</u> | 66.848 | 63.344 | 66.41 | **70.885** |
| Average Mouse TF | 56.434 | 67.996 | <u>71.216</u> | 67.02 | 69.796 | **77.368** |
| Average Core Promoter | 71.81 | 70.527 | 71.07 | 71.627 | <u>75.751</u> | **76.399** |
| Average Promoter_300 | 81.697 | 84.21 | 83.813 | 88.147 | <u>91.15</u> | **92.53** |
| Average Histone | 49.101 | 55.98 | <u>58.832</u> | 58.06 | / | **64.009** |
| Average Overall | 60.69 | 66.649 | <u>67.749</u> | 66.707 | / | **73.334** |

**Table 3:** Nucleotide Transformer Benchmarks (MCC) [10].

| Task | HyenaDNA<br>1.8M | DB<br>89M | DB-2<br>117M | NT<br>2.5B | NT v2<br>500M | AIDO.DNA<br>300M | AIDO.DNA<br>7B |
| --- | --- | --- | --- | --- | --- | --- | --- |
| Enhancer | 52.059 | 49.559 | 52.5 | 54.559 | 55.9 | 49.47 | **59.670** |
| E types | 40.294 | 36.618 | 42.206 | 44.265 | 43.8 | <u>45.067</u> | **45.113** |
| H3 | 78.088 | 76.324 | 78.529 | 79.265 | 78.6 | <u>79.929</u> | **83.207** |
| H3K4me1 | 51.176 | 39.706 | 51.324 | 54.118 | 55 | <u>56.352</u> | **57.39** |
| H3K4me2 | **45.588** | 28.382 | 33.382 | 32.5 | 32 | 36.295 | <u>43.229</u> |
| H3K4me3 | <u>55</u> | 25.882 | 35.294 | 40.882 | 40.6 | 42.841 | **57.42** |
| H3K9ac | <u>58.529</u> | 50.588 | 54.559 | 54.559 | 56.7 | 57.808 | **58.845** |
| H3K14ac | <u>60.882</u> | 40.441 | 51.618 | 53.824 | 54.9 | 56.727 | **65.296** |
| H3K36me3 | 61.324 | 47.353 | 59.118 | 61.765 | 62.4 | <u>64.739</u> | **66.499** |
| H3K79me3 | **66.765** | 57.794 | 61.471 | 62.206 | 63 | 61.911 | 64.156 |
| H4 | 76.324 | 78.382 | 79.706 | 80.735 | 79.9 | <u>82.883</u> | **83.549** |
| H4ac | **56.324** | 35.882 | 46.471 | 49.118 | 49.6 | 53.103 | <u>55.387</u> |
| Promoter all | 91.912 | 92.206 | 94.265 | 95 | **97.6** | 94.355 | <u>95.080</u> |
| Prom no TATA | 91.912 | 92.353 | 94.265 | <u>95.294</u> | **97.6** | 94.39 | 95.244 |
| Prom TATA | 87.941 | 91.029 | 90.882 | 91.765 | **96.5** | 90.801 | <u>92.308</u> |
| Splice Acceptor | 91.618 | x | 94.853 | 97.206 | **98.1** | 98.038 | 97.144 |
| Splice Donor | 89.412 | x | 92.5 | 97.353 | **98.7** | 97.142 | <u>98.216</u> |
| Splice All | 93.382 | 96.176 | 90.882 | 97.206 | **98.4** | 97.657 | <u>97.949</u> |
| Average MCC | 69.363 | 58.667 | 66.879 | 68.979 | 69.961 | <u>69.973</u> | **72.83** |

**Table 4:** Zero-shot variant effect prediction using pretrained model embeddings.

|  | NT [10]<br>2.5B | AIDO.DNA<br>300M | AIDO.DNA<br>7B |
| --- | --- | --- | --- |
| AUROC | .720 | **.807** | .785 |

**Table 5:**
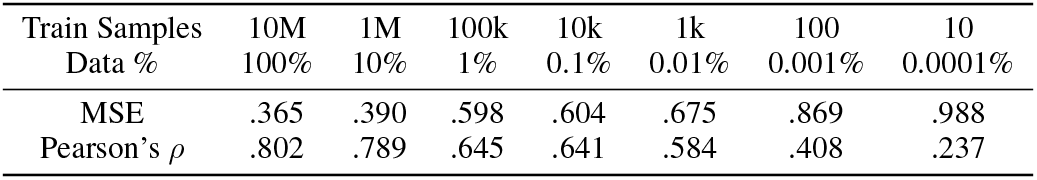
Predicting the activity of promoter sequences with decreasing training data volume, leveraging AIDO.DNA for transfer learning with data scarcity.

**Table 6:**
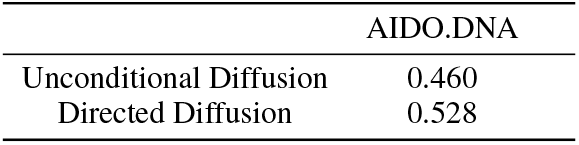
Accuracy of generated promoter sequences.

### 3.2 Scaling Improves Transfer Learning

AIDO.DNA enables high-accuracy recognition of functional genomic elements, revealing a new state-of-the-art on standard benchmarks covering human, mouse, and yeast genomics (Tables 2, 3). These benchmarks, proposed in [10, 11, 4, 7], primarily focus on transcriptional regulation and transcript processing in eukaryotes, which have complicated and sparsely-characterized regulatory grammars. In transfer learning, these accuracy improvements are driven by pretrained representations of DNA function, enabling data-efficient genome mining and experiment prioritization.

### 3.3 Fine-grained Tokenization Enables Accurate Variant Effect Prediction

AIDO.DNA’s single nucleotide tokenization improves recognition of variants with outsize effects on human health, establishing a new SOTA for single-sequence pretrained models (Table 4). While the 300M and 7B versions both exceed the previous SOTA by a wide margin, the smaller 300M model is most performant on this task. We reason that this is due to the smaller embedding size in the 300M model, while the 7B model may have more channels unrelated to variant pathogenicity, creating a lower signal-to-noise ratio on this task.

### 3.4 AIDO.DNA Learns Co-evolving Genome Elements Without Supervision

AIDO.DNA learns functional genomic elements and their interactions without labeled data (Figure 3). Self-supervised pretraining is a powerful tool to infer conditional dependencies, which can be probed and cataloged through *in silico* mutagenesis. While *in silico* mutagenesis studies normally require *O*(*L*^2^) inferences using a supervised model such as Enformer [5] to compute all 2nd order mutation effects, self-supervised models infer the probability of all key mutations under a given query mutation at once, allowing us to compute this dependency mapping with only *O*(*L*) inferences.

Applying dependency mapping to AIDO.DNA reveals a rich landscape of biomolecular elements and interactions (Figure 3). Despite pretraining solely on DNA, we find that AIDO.DNA learns RNA and protein molecules, as well as DNA-protein binding sites, transcriptional regulatory elements, transcription initiation complexes, post-transcriptional modifications, and even predicted RNA secondary structures. Many of these elements are revealed only at the largest 7B model scale.

In one example (Figure 3 left), we apply dependency mapping to the 3’ untranslated region and coding region of KIF26B. While originally known to contain a promoter and coding region, AIDO.DNA also reveals a putative minisatellite enhancer and hairpin-like regulator. Such minisatellites and hairpin-like regions have been implicated in regulation, but exact mechanisms are unknown. The ability of AIDO.DNA to predict functional conservation and co-evolution from a single sequence provides a strong foundation for identifying new genomic functions.

### 3.5 Pretraining Enables Sample-efficient Prediction of Expression from Genomic Contexts

AIDO.DNA accurately and efficiently predicts gene expression from synthetic promoter sequences (Table 5). During transfer learning, AIDO.DNA exploits high-level functional features learned during pretraining, such as promoters, enhancers, transcription factor binding sites (Figure 3), to accurately predict gene expression from genomic contexts with limited labeled data. With just 0.01% of the data (1000 samples), AIDO.DNA retains 73% of the performance on the full 10M dataset. Only 0.001% of the data (100 samples), is required to retain 50% of the top performance.

### 3.6 Directed Generation of Regulatory DNA Sequences

AIDO.DNA learns eukaryotic regulatory sequence grammar, enabling tunable generation of promoters toward a desired level of gene expression (Table 6). We invert the task of expression prediction from sequence, using the same dataset of 10 million random TATA promoter sequences with paired expression level measurements to adapt our model for conditional sequence generation [39]. The dataset contains only random synthetic sequences, allowing us to analytically characterize the performance of an optimal unconditional generation method. With a constant 16bp upstream scaffold and 19bp downstream scaffold on either side of the 80bp random sequence and few bp indels at the insertion sites, a perfect unconditional generator will achieve accuracy of 0.45-0.48. Values above this range require learning promoter grammars which correspond to specific expression levels. Our method, which adapts the pretrained AIDO.DNA with a lightweight 4.5M parameters conditional diffusion head and a simple masked diffusion language modeling objective[38], exceeds this crucial threshold, achieving 0.528 accuracy on per-base recovery when generating sequences *de novo*, using only the desired expression level condition (Table 6).

Alignment and visual inspection of generated sequences under the highest and lowest expression conditions reveals biologically meaningful motifs (Fig. 4). The promoters generated for the highest expression use almost exclusively G to connect the promoter scaffolds with several regions showing no deviation. Those generated for the lowest expression show notable G enrichment at the 5’ end and a noisy alternating TATA motif throughout, possibly hindering expression by re-binding the polymerase after it is recruited to the TATA binding site in the 5’ scaffold.

**Figure 4.**
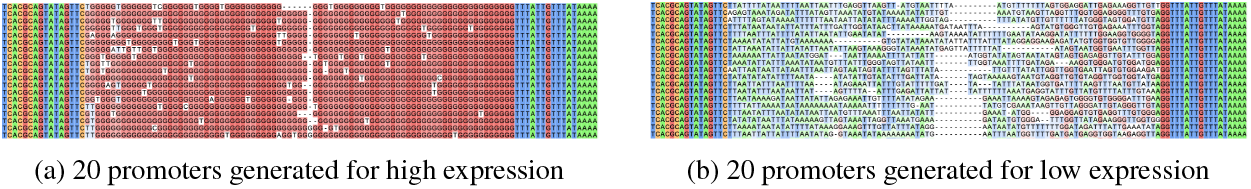
Directed generation of promoter sequences toward a desired expression level. Sequences are aligned and colored by nucleotide with Clustal Omega nucleotide to highlight similarities.

## 4 Discussion

The large feature space of genomes coupled with the relative scarcity of labeled data presents a difficult challenge for statistical modeling approaches, usually dubbed the *big p small n* regime. In this setting, the low information density of the feature space coupled with extreme sample scarcity requires high-bias-low-variance modeling approaches such as additivity or sparsity, or otherwise requires the use of prior knowledge or hand-picked feature sets, none of which are ideal for data-driven research. However, this issue is directly related to a pervasive one-model-one-task approach to research, where users apply curated task-specific data toward a single modeling objective, usually within the scope of a single organism or a single genome. This approach has the dual consequences of limiting the amount of data available for modeling, while also making learned features and effects unlikely to generalize at test-time to new contexts. Especially as heterogeneous and observational data have become more abundant, the one-model-one-task approach seems poorly suited for modern biology research.

Foundation models hint at a new generation of models beyond *big p small n* constraints. Rather than requiring models to learn higher order features and their effects under sample constraints, pretrained foundation models frontload semantic feature learning, and unsupervised models do this without paired or labeled data. To enabling accurate biological models with *big p small n* data, we develop AIDO.DNA, the largest DNA encoder to date with seven billion parameters, which learns informative and transferable representations of DNA by pretraining at scale. We evaluate AIDO.DNA using a wide range of biologically and medically relevant tasks including genome mining and annotation (Tables 2, 3, Fig. 2), synthetic biology (Table 5), therapeutics design (Fig. 4), and disease diagnosis (Table 4). Indeed, we find that unsupervised pretraining learns surprisingly rich representations of DNA which has been mostly unexplored in previous works and only emerge with billions of parameters (Fig. 3).

Despite gaps in performance and utilization relative to protein language models, genome language models are essential for the advancement of biological models beyond the one-model-one-task setting. Genomes are a universal context for biological and medical models, and a recent profusion of context-adaptive modeling methods are well-positioned to take advantage of improved genome representations by contextualizing disease, cell, and patient models with genomic information [40, 41, 42, 43, 44, 45, 46, 47, 48]. AIDO.DNA marks a significant step toward relaxing restrictive model designs that are currently required for *big p small n* data, promising to enable accurate and personalized machine learning in biology and medicine, as well as more general and complex modeling approaches in the development of an AI-driven Digital Organism [1].

## A Data and Code Availability

## B Data and Code Availability

We developed the ModelGenerator package to reproduce, apply, and extend the results in this manuscript https://github.com/genbio-ai/ModelGenerator.

Pre-trained models and finetuning data are also available on Huggingface at https://huggingface.co/genbio-ai.

## C Pre-training Data

Our pre-training dataset is based on the original version of the multi-species dataset from [10].

Our only modifications were to remove genomes which have since been deleted from NCBI, and to replace the deleted *Rattus norvegicus* genome with its current reference genome. We used the included dataloader in this repository, modified to chunk and save non-overlapping 4kbp sequences.

Notably, we found that in the splits from [10] were highly correlated with taxonomy, with the vast majority of the training data being comprised of bacteria, fungi, and invertebrates. In preliminary experiments on reshuffled splits where training included mammals, vertebrates, and popular model organisms, we saw a dramatic decrease in performance as more eukaryotes were added to the training data. While outside the scope of this study, we hypothesize that low-quality, noisy data, such as those common in intergenic eukaryotic sequences, are particularly detrimental to unsupervised model performance. This aligns with the shift in natural language processing to small, highly curated datasets [49]. Although prokaryotes make up the majority of the training set and have drastically different genomes, these genomes have relatively little noise or junk. We reason it may be helpful for DNA language models to learn the basic dependency structures of conservation and co-conservation on these highly constrained genomes, improving performance on eukaryotic data despite the dramatic distribution shift.

## D Pre-training Configs

### D.1 AIDO.DNA 7B

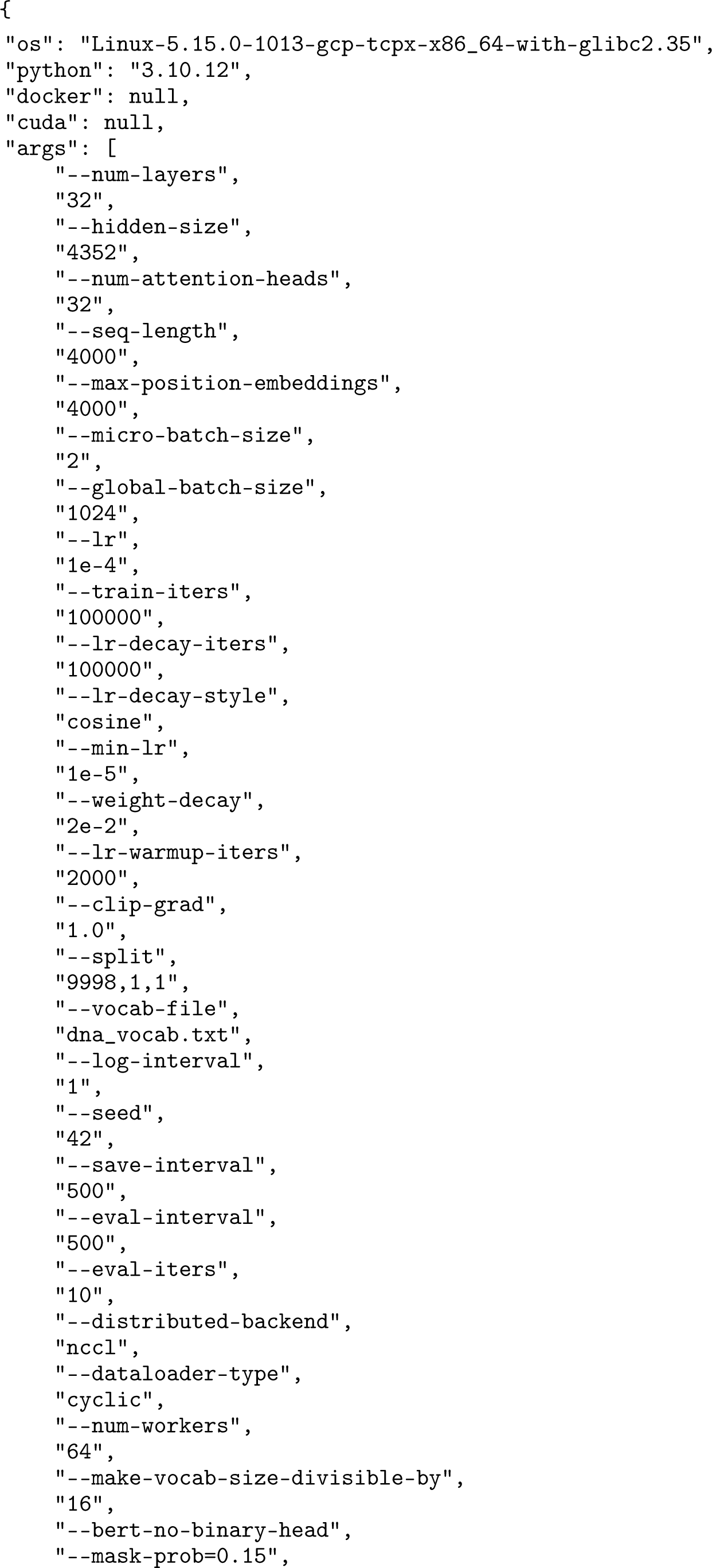
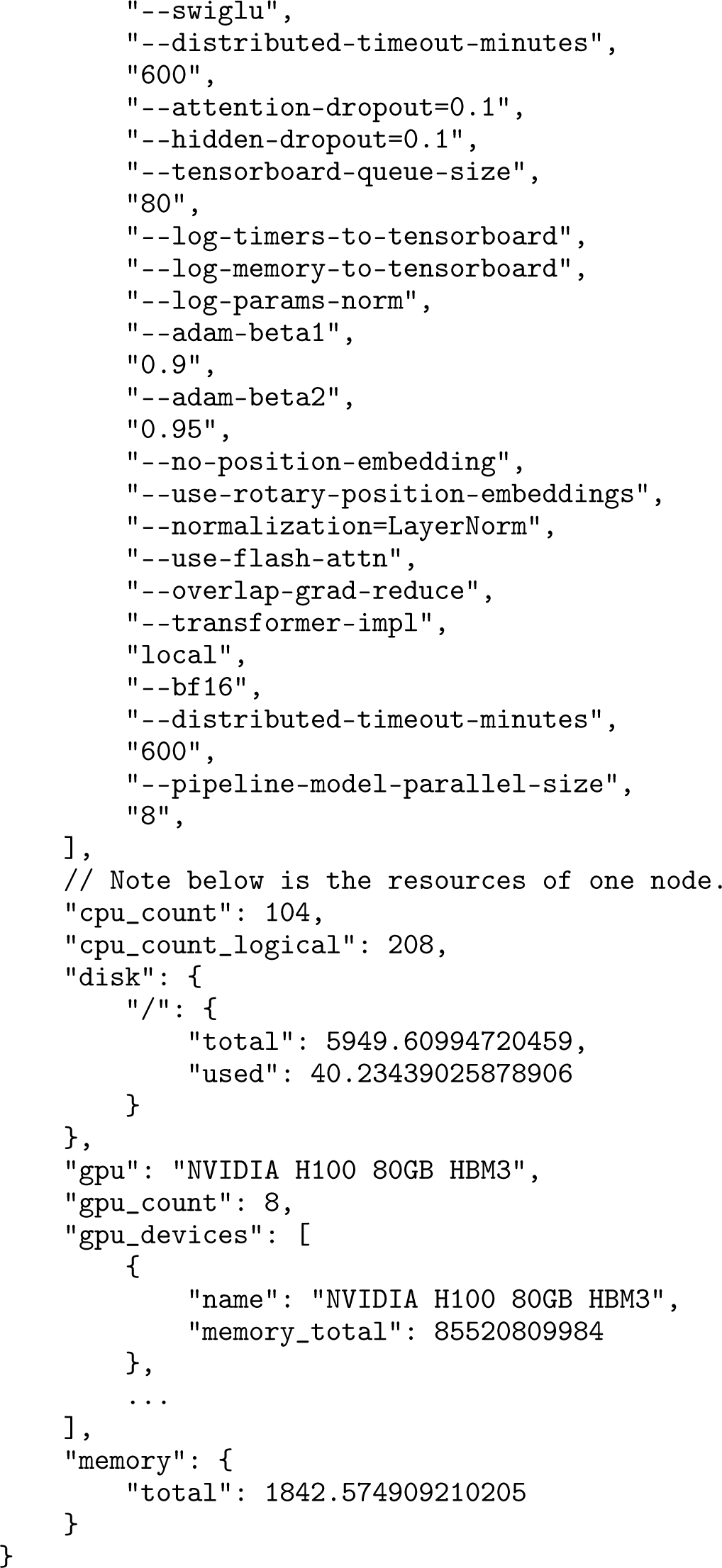

### D.2 AIDO.DNA 300M

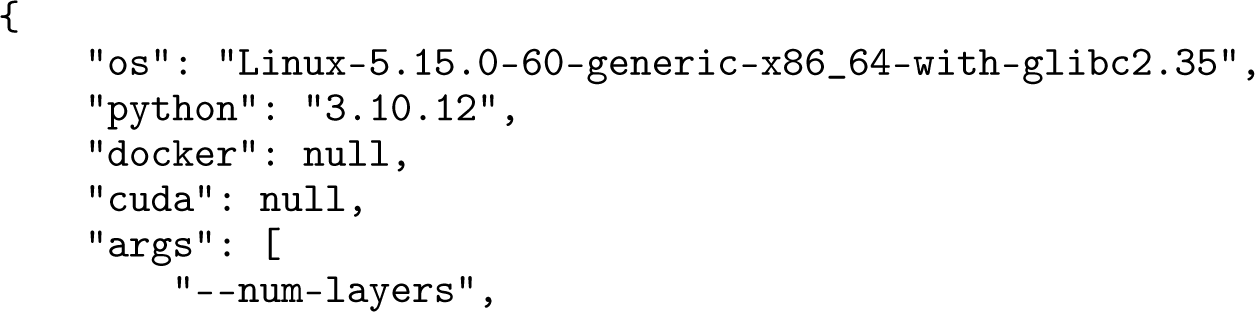
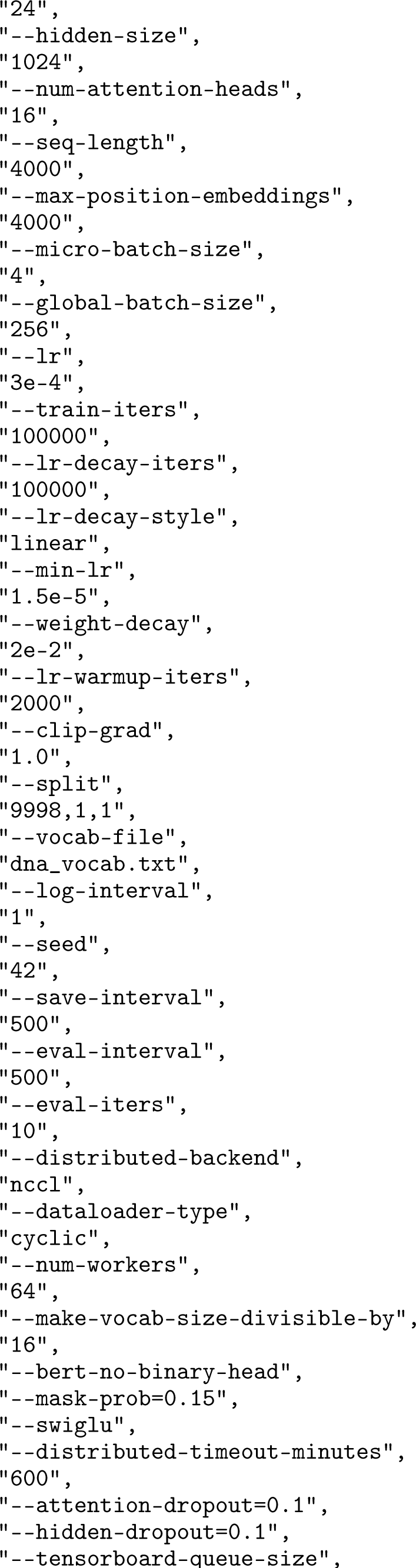
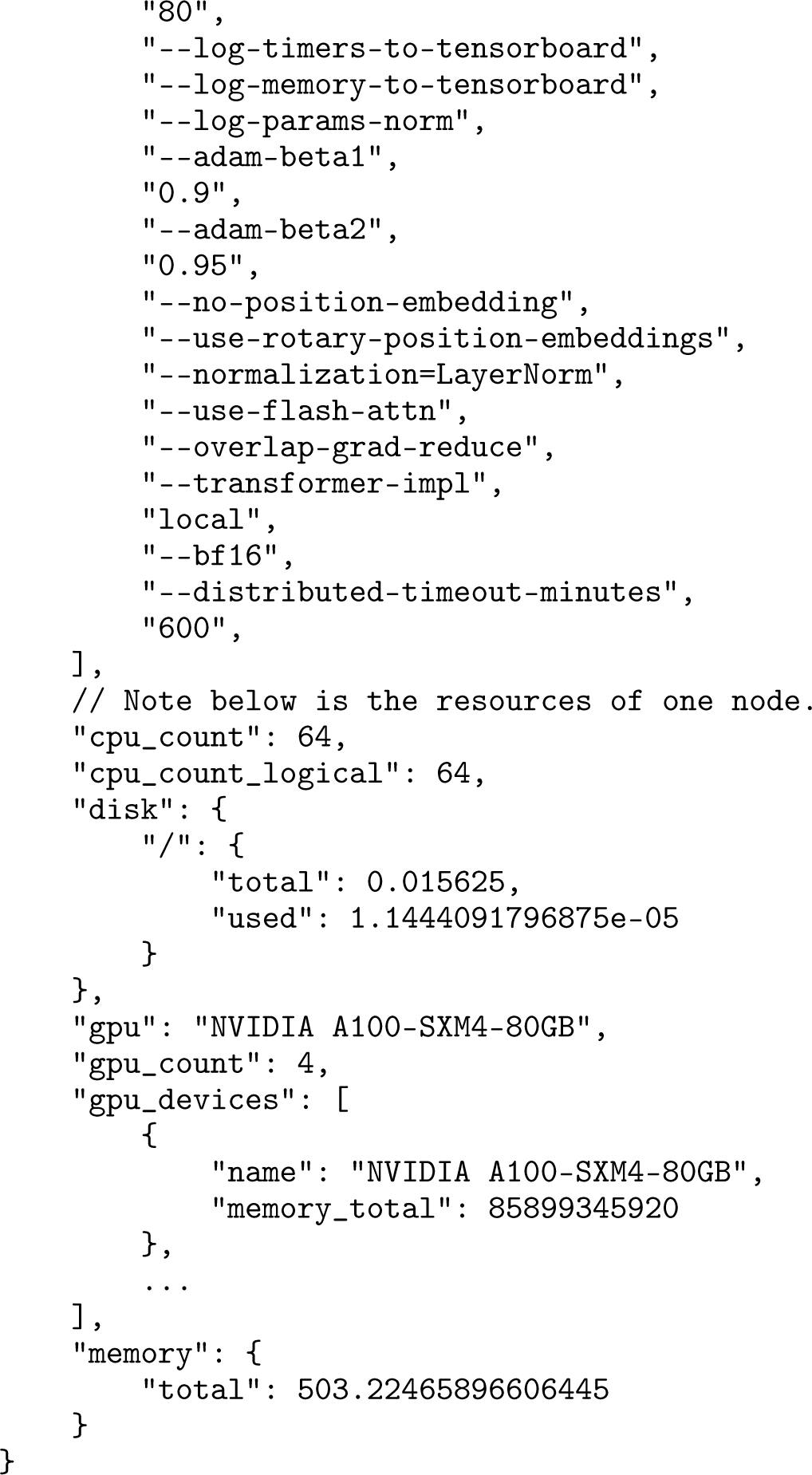

